# HDP-Align: Hierarchical Dirichlet Process Clustering for Multiple Peak Alignment of Liquid Chromatography Mass Spectrometry Data

**DOI:** 10.1101/074831

**Authors:** Joe Wandy, Rónán Daly, Simon Rogers

## Abstract

Matching peak features across multiple LC-MS runs (alignment) is an integral part of all LC-MS data processing pipelines. Alignment is challenging due to variations in the retention time of peak features across runs and the large number of peak features produced by a single compound in the analyte. In this paper, we propose a Bayesian non-parametric model that aligns peaks via a hierarchical cluster model using both peak mass and retention time. Crucially, this method provides confidence values in the form of posterior probabilities allowing the user to distinguish between aligned peaksets of high and low confidence. The results from our experiments on a diverse set of proteomic, glycomic and metabolomic data show that the proposed model is able to produce alignment results competitive to other widely-used benchmark methods, while at the same time, provide a probabilistic measure of confidence in the alignment results, thus allowing the possibility to trade precision and recall.

**Availability:** Our method has been implemented as a stand-alone application in Java, available for download at http://github.com/joewandy/HDP-Align.

## 1 Introduction

Liquid-chromatography coupled to mass-spectrometry (LC-MS) is a popular method for performing large-scale experiments and investigating the differential expression of compounds in LC-MS-based-omics (such as proteomics, metabolomics and glycomics). Large-scale untargeted LC-MS studies may involve the analyses of potentially hundreds of runs [6]. Typical LC-MS data pre-processing pipelines operate in a serial manner with many intermediate steps. In untargeted LC-MS studies, the presence of even relatively small errors in any steps preceding the identification stage could potentially result in significant differences to the final analysis and biological conclusions. Alignment, where the correspondences between peaks across runs are established, forms an integral part of the LC-MS data preprocessing pipeline and is a challenging problem due to the non-linear deviations in retention time (RT) of peak features across runs [23] and the large number of ‘related-peaks’ derived from a single compound alone [9]. These related-peaks include e.g. adducts, fragments and multiple ionisation forms, and often share similar chromatographic peak shape correlations and close RT values. In this paper, the term ‘run’ refers to the output from running any biological sample through the LC/MS instrument once, while the term ‘feature’ refers to a tuple of minimally the (*m/z*, *RT*) values derived from a single peak.

Alignment methods can be broadly divided into two categories [27]: (a) warping-based methods that perform RT correction of peak features before matching, and (b) direct-matching methods that establish peak correspondence by performing matching on peak features directly, without performing RT correction, often by maximising some objective function. Warping methods attempt to correct the RT drifts present across runs, either using the full LC-MS profile data or the feature data alone. Once the time warping function has been established, finding the actual matching between peak features is straightforward as both peak masses and RT values should be accurately conserved across runs. In contrast, direct matching methods perform the alignment of peaks by directly matching the features across runs, skipping the initial time-warping step. Direct matching methods can be preferred in certain cases due to their simplicity, while still offering good performance [14,22,33].

Alignment methods can also be categorised depending on whether they require a user-defined reference run to be specified. When such reference is necessary, the full alignment of multiple runs is constructed through successive merging of pairwise runs towards the reference run (e.g. MZmine2’s Join aligner in [22]. Alternatively, methods that do not require a reference run can either operate in a hierarchical fashion – where the final multiple alignment results are constructed in a greedy manner by merging of successive pairwise results following a guide tree (e.g. SIMA, described in [33]) – or by pooling features across runs and grouping similar peaks in the combined input simultaneously (e.g. the *group()* function of XCMS in [26]).

According to [27], the common shortcomings shared by many alignment methods include the incorrect modelling assumption that elution order of peaks is preserved across runs and the abundance of user-defined parameters, which can dramatically influence alignment results. Further uncertainties can be introduced due to the selection of a reference run and the construction of a guide tree in hierarchical methods. Since alignment is such an important part of the data preprocessing steps, it would be useful to be able to robustly identify the uncertainty or confidence in the alignment results. The subject of identifying and quantifying uncertainty has been extensively investigated in the problem of multiple sequence alignment (MSA) for genomics and transcriptomics. [13] attempt to quantify the alignment uncertainty of the popular MSA tool ClustalW [30], based on evaluations using synthetic data, and concludes that between half to all columns in their benchmark MSA results contain alignment errors. [19] construct a score that reflects the consensus between all possible pairwise alignments in T-COFFEE, while [20] propose GUIDANCE, a confidence measure obtained from pertubations of guide trees. Statistical approaches that provide a measure of confidence in alignment results have also been explored by [24] and [2], where the MSA results and phylogeny are constructed simultaneously, thus eliminating the need for a guide tree.

Despite the clear benefits of alignment uncertainty quantification in the sequence domain, the challenge of quantifying alignment results remains relatively unaddressed for the alignment of multiple runs in LC-MS-based-omics. Bayesian methods operating on profile data [11,16,32] and feature-based alignment methods [7,22, 33] exist to correct RT drift, but in such methods, uncertainties are not propagated from the RT regression stage to the necessary peak matching stage that follows. Several recent feature-based alignment methods incorporate probabilistic modelling as part of their workflow, making it possible to extract some form of scores or probabilities on the alignment results. These methods are often limited to the alignment of two runs, which is not a realistic assumption in actual LC-MS experiments. For example, [10] propose an empirical Bayes model for pairwise peak matching. Matching confidence can be obtained from the model in form of posterior probability for any peak pair in two runs, however constructing multiple alignment results in [10] still requires a greedy search to find candidate features within m/z and RT-RT tolerances to a predetermined set of ‘landmark’ peaks. [8] describe PeakLink, a workflow for alignment that performs an initial warping using a fourth-degree polynomial. PeakLink poses the pairwise matching problem as a binary classification problem, where a Support Vector Machine (SVM) is trained based on an alignment ground truth derived from MS-MS information and used to differentiate matching and non-matching candidate pairs to produce the actual alignment results. While not explicitly included in the output of PeakLink, a matching score can be extracted from the SVM that represents how far each candidate pair is from the decision boundary. Note that these scores are not well-calibrated in the probabilistic sense, thus making comparisons of matching scores less straightforward. PeakLink is also not extended to the problem of aligning multiple runs, although [8] state that it would be possible to do so with the choice of a suitable reference run.

The goal of establishing the matching of peaks across multiple runs at once can be viewed as a clustering problem, where a set of peaks can be grouped (by their m/z, RT and other suitable features) into local clusters within each run (representing all of the peaks from an individual compound), which are further grouped into global clusters shared across runs. A preliminary form of this idea has been explored in [5], where hierarchical clustering is performed on the total ion chromatogram data to group peaks into within-run local clusters, which are further grouped into across-run super clusters. The highly accurate mass information available from modern LC-MS platforms is not used in [5], although it is highlighted as a possible future work. The choice of using a hierarchical clustering method in [5] also requires choosing various user-defined parameters, such as determining a suitable cut-off for the dendogram produced, deciding on a suitable linkage method and defining an appropriate distance measure between groups of peaks.

In this work, we expand upon the idea of viewing direct matching as a hierarchical clustering problem by proposing **HDP-Align**, a Bayesian non-parametric model that groups related-peaks within runs and assigns them to global clusters shared across runs. Within each global cluster, peaks are further grouped by their m/z values into mass clusters, which represent the various ionisation products (IPs) derived from the global compound. The proposed model allows us to infer the matching of peaks across all runs at once, without the need for any intermediate merging of pairwise runs, and the resulting posterior summaries provide us with a confidence score in the matching quality of aligned peaksets. This introduces the possibility of allowing the user to trade recall for precision from the alignment results by returning a smaller subset of the results having a higher confidence score of being correctly aligned. Figure 1 shows an illustration of the clustering process in HDP-Align. Additionally, the latent variables of clustering structure inferred in the model can potentially have physically meaningful identities that can be used for further analysis, and using a metabolomic dataset, we demonstrate the usefulness of such clustering objects by using the mass clusters derived from the model to perform putative annotations of features based on their potential adduct types and metabolite identities.

**Figure 1.**
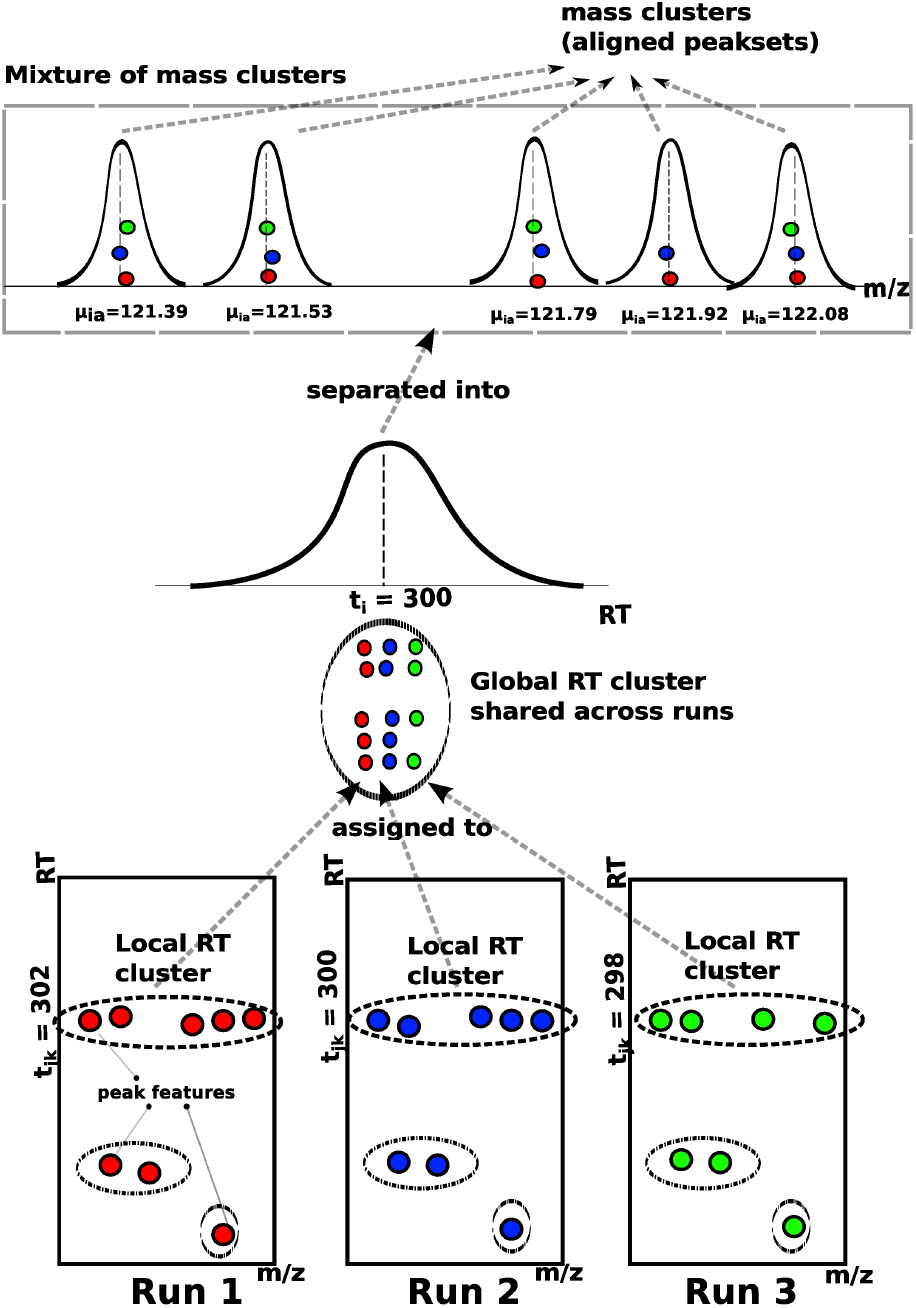
An illustrative example of how the proposed model in HDP-Align simultaneously (**1**) performs the clustering of related peak features into within-run local clusters by their RT values, (**2**) assigns the peak features to global RT clusters shared across runs, and (**3**) separates peak features into mass clusters, which correspond to aligned peaksets.

## 2 Related Work

The goal of establishing the matching of peaks across multiple runs at once can be viewed as a clustering problem, where a set of peaks can be grouped (by their m/z, RT and other suitable features) into local clusters within each run (representing all of the peaks from an individual compound), which are further grouped into global clusters shared across runs. Hierarchical clustering has been used for the matching of peaks across runs [5,31]. In [31], peaks are hierarchically clustered based on their m/z values to construct matching across runs, while in [5], peaks from the entire dataset are pooled and a hierarchical clustering scheme based on RT only is used to group peaks into within-run local clusters, which are further grouped into across-run super clusters. Both approaches require choosing various user-defined parameters, such as determining a suitable cut-off for the dendogram produced, deciding on a suitable linkage method and defining an appropriate distance measure between groups of peaks. In [31], no chromatographic separation is performed, so only the m/z values of peaks are used. The nature of the gas chromatography data used in [5], where retention time across runs is more reproducible, means that even without using the m/z information, good alignment performance can still be obtained. This will not be the case of LC-MS data, where retention time drift is common and the highly accurate m/z information is crucial for alignment. The proposed HDP-Align model fills this gap where both m/z and RT values, important for LC-MS peak alignment, are used for the hierarchical clustering process. The probabilistic approach employed by HDP-Align also allows us to extract confidence values from aligned peaksets.

## 3 Hierarchical Dirichlet Process Mixture Model for Alignment

The proposed model for HDP-Align is framed as a Hierarchical Dirichlet Process (HDP) mixture model [29]. Essential modifications to the basic HDP model were performed to suit the nature of the multiple peak alignment problem. Figure 2 shows the conditional dependencies between random variables in the HDP-Align model.

**Figure 2.**
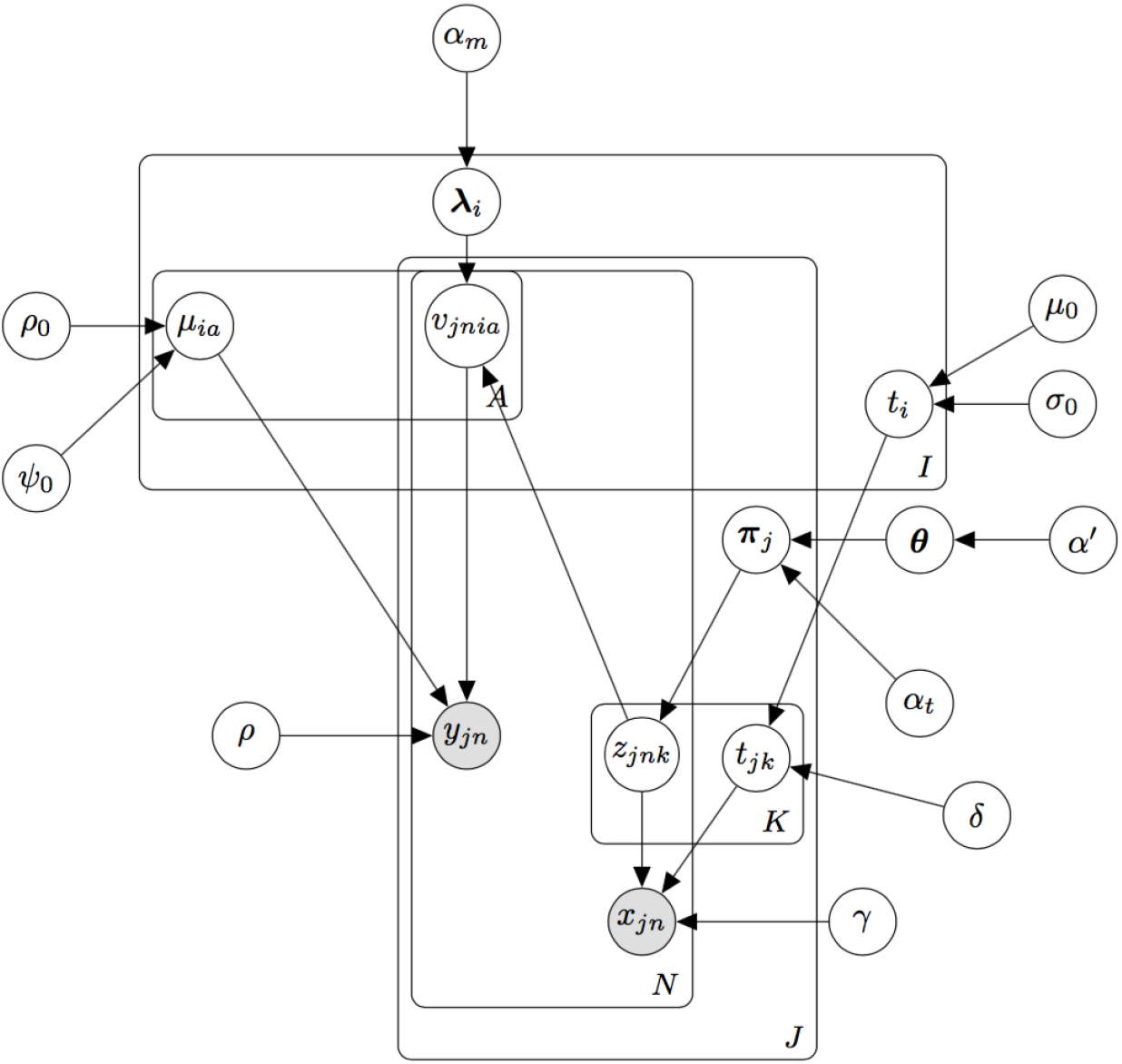
Graphical model for HDP-Align. *x*_*jn*_ is the observed RT value of peak *n* in file *j*, while *y*_*jn*_ is the observed m/z value.

Our input consists of *J* input files, indexed by *j* = 1, …, *J*, corresponding to the *J* LC-MS runs to be aligned. Each *j*-th input file contains *N*_*j*_ peaks in total, which can be separated into *K*_*j*_ local clusters of related-peaks. In a *j*-th file, peaks are indexed by *n* = 1, …, *N*_*j*_ and local clusters are indexed by *k* = 1, …, *K*_*j*_. Across all files, we assign each local cluster *k* in file *j* to a global cluster *i* = 1, …, *I*, where *I* is the total number of global clusters, using the indicator variable *v*, as described in the following paragraph. A global cluster corresponds to the compound of interest during LC-MS analysis, e.g. metabolite or peptide fragment, that is present across runs, while local clusters are realisations of the global clusters in a specific run. Finally, within each global cluster *i*, we can further group peaks by their m/z values into A mass clusters (indexed by *a* = 1, …, *A*). Each mass cluster therefore corresponds to the ionization product peaks coming from the different runs that are produced by a global compound during mass spectrometry.

We use the indicator variable *z*_*jnk*_ = 1 to denote the assignment of peak *n* in file *j* to local cluster *k* in that file. Similarly, *v*_*jni*_ = 1 if peak *n* in file *j* is assigned to global cluster *i*, and *v*_*jnia*_ = 1 if peak *n* in file *j* is assigned to mass cluster *a* linked to metabolite *i*. Let *d*_*j*_ be the list of observed data of peaks in file *j*, *d*_*j*_ = (**d**_*j*1_, **d**_*j*2_, …, **d**_*jn*_) where **d**_*jn*_ = (*x*_*jn*_, *y*_*jn*_) with *x*_*jn*_ the RT value and *y*_*jn*_ the log m/z value of the peak feature. The log of m/z value is here used as the m/z error is assumed to increase linearly with the observed m/z value [21]. ***θ*** denotes the global mixing proportions and *π*_*j*_ the local mixing proportions for file *j*. The global mixing proportions ***θ*** are distributed according to the Griffiths, Engen and McCloskey (GEM) distribution:

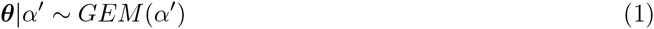

where the *GEM* distribution over ***θ*** is described through the stick-breaking construction:

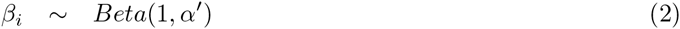

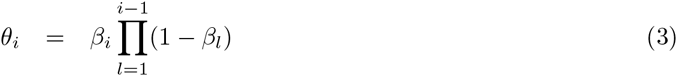

The local mixing proportions *π*_*j*_ are distributed according to a Dirichlet Process (DP) prior with the base measure ***θ*** and concentration parameter *α*_*t*_.

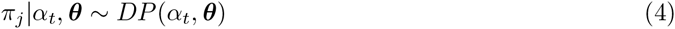

Within each file *j*, the indicator variable *z*_*jnk*_ = 1 denotes the assignment of peak *n* in file *j* to local RT cluster *k* in that file. This follows the local mixing proportions for that file.

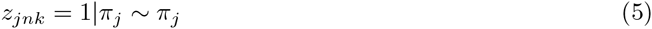

The RT value *t*_*i*_ of a global mixture component is drawn from a base Gaussian distribution with mean *µ*_0_ and precision (inverse variance) *σ*_0_.

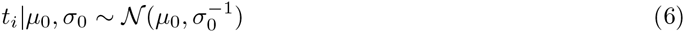

The RT value *t*_*ij*_ of a local mixture component in file *j* is normally distributed with mean *t*_*i*_ and precision *δ.* The precision controls how much RT values of related-peak groups across runs are allowed to deviate from the parent global compound’s RT.

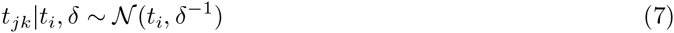

Finally, the observed peak RT value is normally distributed with mean *t*_*jk*_ and precision γ. The precision controls how much RT values of peaks can deviate from their related-peak group.

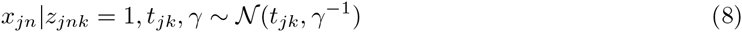

The m/z value produced through high-precision MS instrument is highly accurate, and its correspondence is often preserved across runs. Once peaks have been assigned to their respective global clusters, we need to further separate peaks within each global cluster into mass clusters to obtain the actual alignment. These mass cluster corresponds to ionisation products. We do this by incorporating an internal DP mixture model on the m/z values (*y*_*jn*_) within each global cluster *i.* Let the indicator *v*_*jnia*_ = 1 denotes the assignment of peak *n* in file *j* to mass cluster *a* in the *i*-th global cluster. Then:

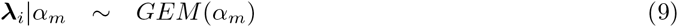

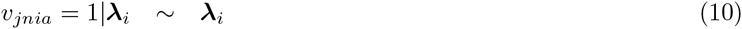

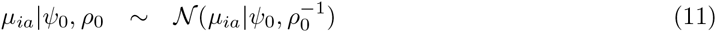

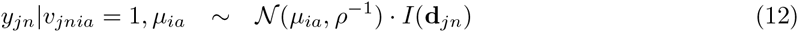

where the index *ia* refers to the *a*-th mass cluster of the *i*-th global cluster. **λ**_*i*_ is the mixing proportions of the *i*-th internal DP mixture for the masses, with *α*_*m*_ the concentration parameter. *µ*_*ia*_ is the mass cluster mean, drawn from the Gaussian base distribution with mean *ψ*_0_ and precision *ρ*_0_. The observed mass value is drawn from a Gaussian distribution with the component mean *µ*_*ia*_ and precision *ρ*, for which the value is set based on the MS instrument’s resolution. Additionally, we add an additional constraint on the likelihood of *y*_*jn*_ using the indicator function *I*(·) such that *I*(**d**_*jn*_) = 1 if there are no other peaks inside the mass cluster that come from the same file as the current **d**_*jn*_ peak, and 0 otherwise. This constraint captures the restriction that a peak feature can only be matched to other peaks from different files, reflecting the assumption that within each LC-MS run, compounds produce ionisation products with distinct mass-to-charge fingerprints that can be used for matching to other runs.

## 4 Inference

Inference within the model is performed via a Gibbs sampling scheme, allowing us to compute the alignment probabilities through the proportion of posterior samples in which any sets of peaks are placed in the same mass component (*a*) in the same top-level cluster. In this manner, peaks coming from different runs that are in the same mass component are considered to be aligned as they have similar RT and m/z values. In each iteration of the sampling procedure, we instantiate the mixture component parameters for the local RT cluster (*t*_*jk*_) and global RT cluster (*t*_*i*_) in the mixture model. In the internal DP mixture linked to each global cluster *i*, we marginalise out the mass cluster parameters (*µ*_*ia*_). The initialisation step of the sampler is performed by assigning all peaks in each run into a single local RT cluster. Across runs, these local clusters are assigned under a global cluster shared across runs. Within a global cluster, peaks coming from different runs are assigned to a single mass cluster. The sampler than iterates through each peak feature, removing it from the model, updating the assignment of peaks to clusters and performing the necessary book-keeping on any instantiated mixture components. Further details on the specific Gibbs update statements can be found in following sections.

### 4.1 Updating peak assignments

We use the following variables to denote the count of items in any clustering object: *c*_*jk*_ is the number of peaks in a local cluster *k* of file *j. c*_*i*_ is the number of local clusters in a global cluster *i*, and *c*_*ia*_ is the number of peaks in a mass cluster *a* inside a global RT cluster *i.* To update the assignment of a peak **d**_*jn*_ to local RT cluster *k* during Gibbs sampling, we need the conditional probability of *P*(*z*_*jnk*_ = 1) given every other parameters, denoted as *P*(*z*_*jnk*_ = 1|**d**_*jn*_,…).

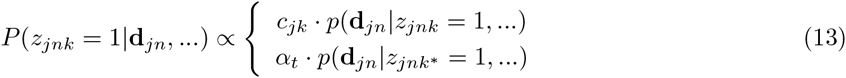

We consider the top and bottom terms of eq. (13) separately in the following.

1. The likelihood of the peak **d**_*jn*_ to be in an existing local RT cluster *k, p*(**d**_*jn*_|*z*_*jnk*_ = 1,…) is proportional to *c*_*jk*_. This is assumed to factorise across the RT (*x*_*jn*_) and mass (*y*_*jn*_) terms

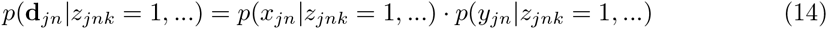 The RT term *p*(*x*_*jn*_|*z*_*jnk*_ = 1,…) in eq. (14) is normally distributed with mean *t*_*jk*_ and precision γ, while the mass term *p*(*y*_*jn*_|*z*_*jnk*_ = 1, …) is an internal DP mixture of mass components linked to the parent global cluster *i* of an existing local cluster *k.* We then marginalise over all mass clusters in *i* to get *p*(*y*_*jn*_|*z*_*jnk*_ = 1, *v*_*jni*_ = 1…)

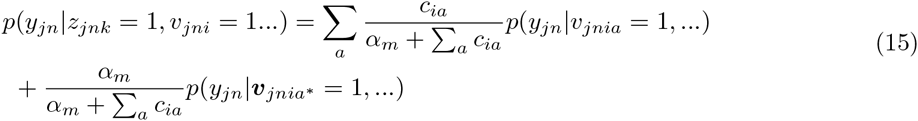 To compute the terms in eq. (15), first we consider the case for an existing mass cluster *a* in the global RT cluster *i.* Then,

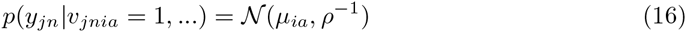 For a new mass cluster *a\** in the global RT cluster *i*, we marginalise out *µ*_*ia*_ to obtain

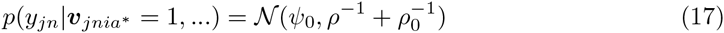
2. The likelihood of the peak **d**_*jn*_ to be in a new local cluster *k\** is proportional to *α*_*t*_. Marginalising over all global clusters in the top-level DP, we get

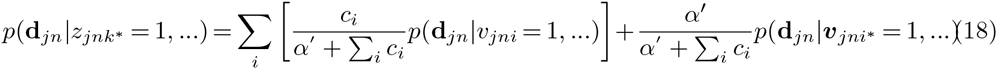 There are two terms to compute in eq. (18): whether peak **d**_*jn*_ is in an existing global cluster *i* or a new global cluster *i*.* For an existing global RT cluster *i* in eq. (18), *p*(**d***j*_*n*_|*v*_*jni*_ = 1,…) is assumed to factorise into its RT and mass terms, so *p*(**d**_*jn*_|*v*_*jni*_ = 1,…) = *p*(*x*_*jn*_|*v*_*jni*_ = 1,…) *· p*(*y*_*jn*_|*v*_*jni*_ = 1, …). We marginalise over all local RT clusters to obtain

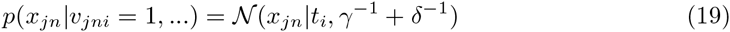

and marginalise over all possible mass clusters in the internal DP linked to global cluster *i* to obtain *p*(*y*_*jn*_|*v*_*jni*_ = 1, …). This is defined in eq. (15). Finally, for a new global RT cluster *i\** in eq. (18), *p*(**d**_*jn*_|***v***_*jni\**_ = 1,…) is also assumed to factorise into its RT and mass terms. Then, we marginalise over *t*_*jk*_ and *t*_*i*_ to obtain

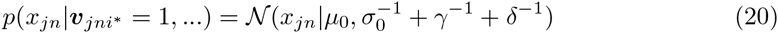

and marginalise over *µ*_*ia*_ to get

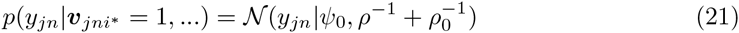

### 4.2 Updating instantiated variables

The following expressions are used to update the instantiated mixture component parameters in the model during Gibbs sampling.

1. Updating global cluster’s RT *t*_*i*_: here, *t*_*jk∊i*_ refers only to local RT clusters currently assigned to the global cluster *i*, and *c*_*i*_ is the count of such peaks. Then

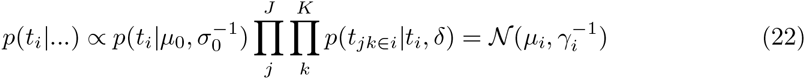

where 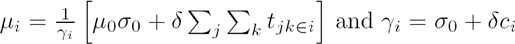.
2. Updating local cluster’s RT *t*_*jk*_: here, *x*_*jn∊k*_ refers only to peaks currently assigned to the local RT cluster *k*, and *c*_*jk*_ is the count of such peaks.

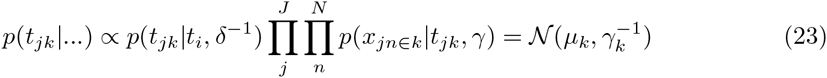

where 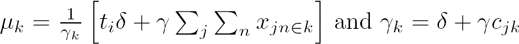.

### 4.3 Using the Inference Results

Using the posterior samples from Gibbs sampling, we can compute various posterior summaries and more interestingly, extract the alignment of peaks from the inference results (since features assigned into the same mass cluster within the same global RT cluster are considered to be aligned). For each sample, we record the aligned peaksets of peaks put into the same mass cluster. Averaging over all samples provides a distribution over these aligned peaksets. Note that across all the aligned peaksets from all samples, it is possible for the same peak to be matched to different partners with varying probabilities, depending on how often they co-occur together in the same mass cluster. To allow the possibility of controlling precision and recall from the results, we provide another user-defined threshold *t*, where aligned peaksets are returned only when they occur with matching probability >*t*. Varying this threshold allows the user to use HDP-Align to trade precision for recall: a low value for *t* gives a larger set of results that are potentially less precise, while conversely a high *t* provides a smaller, more precise set of aligned peaksets. This is an output not available from the other baseline alignment methods and can potentially be useful in problem domains where high precision is required from the alignment results.

### 4.4 Isotopic Product and Metabolite Identity Annotations

In metabolomic studies using electrospray ionisation, a single metabolite can generate multiple ionisation products (IPs, such as isotopic variants, adducts, fragment peaks), alongside other peaks resulting from noise and artifacts introduced during mass spectrometry [15]. Determining and annotating these IP peaks are desirable to remove extraneous peaks and reduce the burden of subsequent analysis in the data processing pipeline. Additionally, deducing the precursor mass of the compound that generates the IP peaks is necessary to query compound library databases in order to assign metabolite identities.

The resulting clustering objects inferred from HDP-Align lend themselves to further analysis in a natural fashion, as global RT clusters in HDP-Align may correspond to metabolites, while local RT clusters may correspond to the noisy realisations of these metabolites within each run. Mass clusters in the internal mixture of each global cluster could correspond to the ionisation products of a metabolite. To demonstrate the possibility of obtaining additional information beyond alignment from the output of HDP-Align, we follow the workflow in [15] that performs IPs and metabolite annotations of peaks. This workflow is composed of multiple key steps: peak matching, ionisation product clustering and metabolite mass matching. A key difference of HDP-Align to the workflow in [15] lies in the fact that HDP-Align is able to perform the two separate steps of peak alignment and potential IP clustering simultaneously, as shown in Figure 3.

**Figure 3.**
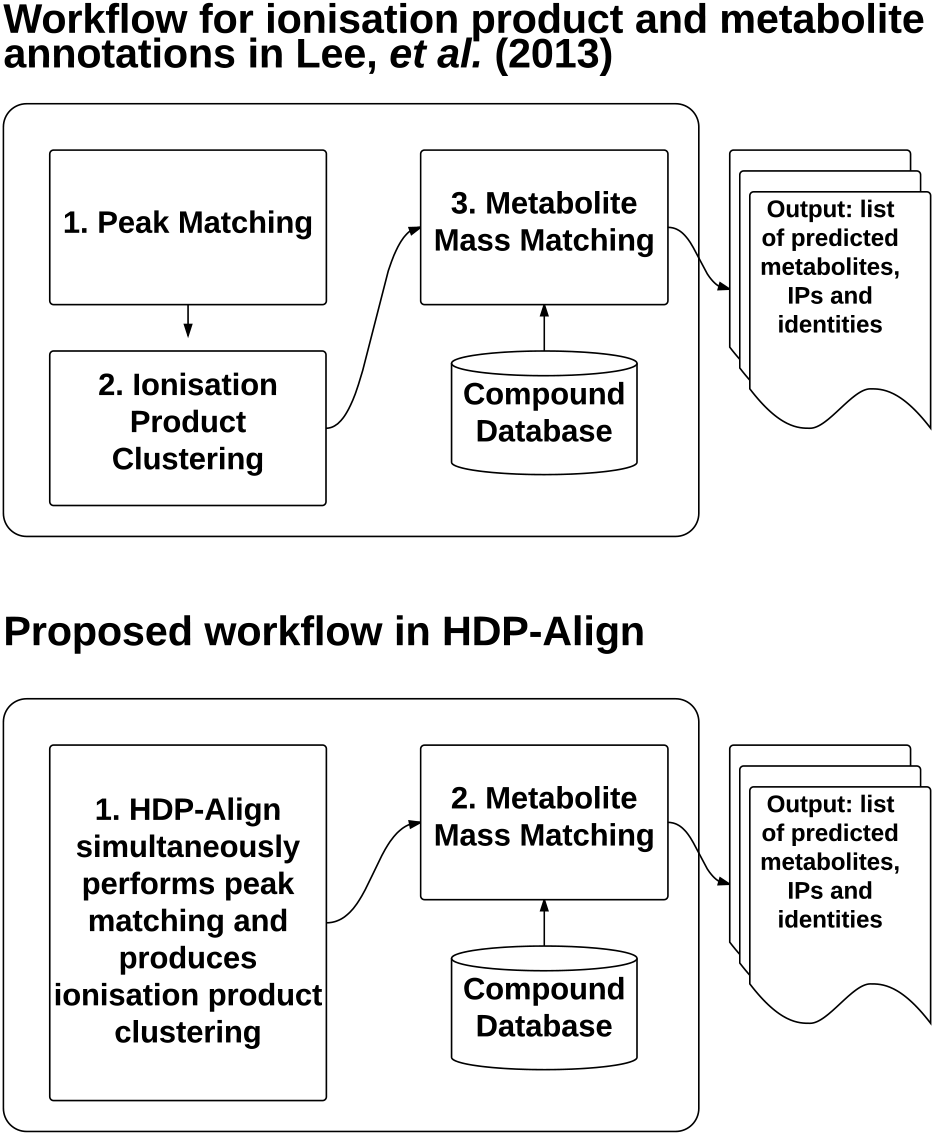
Comparisons on the workflow to assign putative annotations on isotopic products and metabolites described in Lee. *et al*. (2013) [15] and in HDP-Align.

Given the set of potential IP clusters, we can perform IP annotations on the peaks. To do this using the metabolomic dataset, first we take the set of clustering objects produced in a posterior sample. For each mass cluster, we assign its m/z value to be the average m/z values of features assigned to it, denoted by *m*. The list of common adducts (Table 1) in positive ionisation mode is used to compute the inverse transformation for the precursor mass that generates the observed mass. Following [15], any two mass clusters sharing the same precursor mass (within tolerance) provide a vote on the presence of that consensus precursor mass. The mass clusters and peaks inside them can be annotated with the adduct type that produces the transformation type to the shared precursor mass. The set of precursor masses deduced in this manner can also be used to query KEGG (a database of metabolite compounds) in order to assign metabolite identities to the global compound.

## 5 EVALUATION STUDY

### 5.1 Evaluation Datasets

Performance of the proposed methods and other benchmark methods is evaluated on LC-MS datasets from proteomic, glycomic and metabolomic experiments (Table 5 summarises the number of features in the datasets). The Proteomic dataset is obtained from [14]. All 6 fractions from the Proteomic dataset in [14], each containing 2 runs of features having high RT variations across runs, are used in our experiments. The Glycomic dataset is provided by [32]. We use the first 10 runs from the dataset in our experiment. Both of the Proteomic and Glycomic datasets provide alignment ground truth and have been used to benchmark alignment performance in other studies [1,14,22,32,33]. Additionally, we also introduce a metabolomic dataset generated from standard runs used for the calibration of chromatographic columns [3]. The runs were produced from different LC-MS analyses separated by weeks, representing a potentially challenging alignment scenario. 6 runs were used in the experiment. Alignment ground truth was constructed from the putative identification of peaks in each of the 6 runs separately at 3 ppm using the Identify module in mzMatch [25], taking as additional input a database of 104 compounds known to be present and a list of common adducts in positive ionisation mode (Table 1). This is followed by matching of features sharing the same annotations across runs to construct the alignment ground truth. Only peaks unambiguously identified with exactly one annotation are used for this purpose; peaks with more than one annotation per run are discarded from the ground truth construction.

**Table 1.** List of common adduct types in positive ionisation mode for ESI.

| Adduct Types |  |  |  |
| --- | --- | --- | --- |
| M+2H | M+H | M+ACN+H | M+H+NH<sub>4</sub> |
| M+NH<sub>4</sub> | M+ACN+Na | 2M+ACN+H | M+ACN+2H |
| M+Na | M+2ACN+H | M+2ACN+2H | M+CH<sub>3</sub>OH+H |
| 2M+H |  |  |  |

Table 2 summarises the different evaluation datasets and the number of features each dataset has.

**Table 2.**
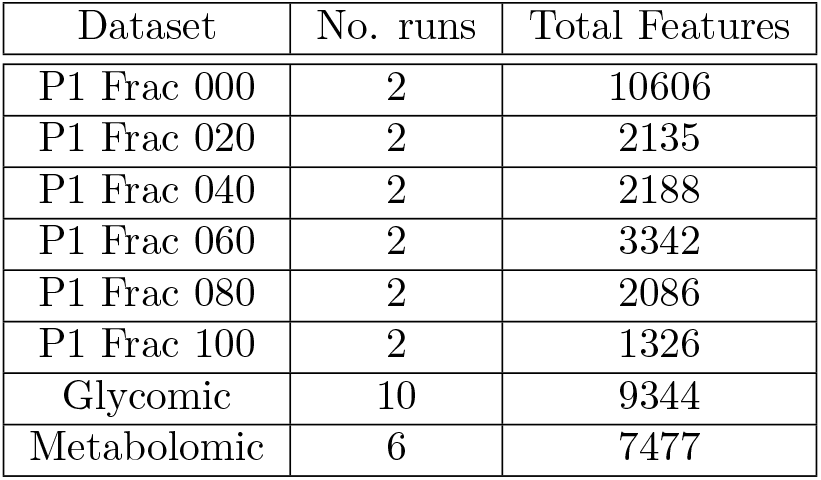
Total number of runs and features of the selected evaluation datasets.

| Dataset | No. runs | Total Features |
| --- | --- | --- |
| P1 Frac 000 | 2 | 10606 |
| P1 Frac 020 | 2 | 2135 |
| P1 Frac 040 | 2 | 2188 |
| P1 Frac 060 | 2 | 3342 |
| P1 Frac 080 | 2 | 2086 |
| P1 Frac 100 | 2 | 1326 |
| Glycomic | 10 | 9344 |
| Metabolomic | 6 | 7477 |

### 5.2 Performance Measures

Performance of the evaluated methods in our results is quantified in terms of precision and recall. These two measures are also commonly used in information retrieval, where ‘precision’ refers the fraction of retrieved items that are relevant, while ‘recall’ refers the fraction of relevant items that are retrieved [18].

To provide a definition of ‘precision’ and ‘recall’ suitable for evaluating alignment performance, we first enumerate all the possible *q*-size combinations for every aligned peakset in both the method’s output and the ground truth list. For example, an alignment method returns a list of two aligned peaksets {*a*, *b*, *c*, *d*,}, {*e*, *f*, *g*} as output. When *q* = 2, this output can be enumerated into a list of 9 ‘alignment items’ of all the pairwise combinations of features: {*a*, *b*}, {*a*, *c*}, {*a*, *d*}, {*b*, *c*}, {*b*, *d*}, {*c*, *d*}, {*e*, *f*}, {*e*, *g*}, {*f*, *g*}. Let *M* and *G* be the results from such enumeration from a method’s output and the ground truth respectively. Each distinct combination of features in *M* and *G* can be considered as an item during performance evaluation. Intuitively, the choice of *q* reflects the strictness of what is considered to be a true positive item, with larger values of *q* demanding an alignment method that produces results spanning more runs correctly.

For a given *q*, the following positive and negative instances of alignment item can now be defined for the purpose of performance evaluation:

- True Positive (***TP***): items that should be aligned (present in *G*) and are aligned (present in *M*).
- False Positive (***FP***): items that should not be aligned (absent from *G*) but are aligned (present in *M*).
- True Negative (***TN***): items that should not be aligned (absent from *G*) and are not aligned (absent from *M*).
- False Negative (***FN***): items that should be aligned (present in *G*) but are not aligned (absent from *M*).

In the context of alignment performance, precision 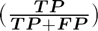 is therefore the fraction of items in *M* that are correct with respect to *G*, while recall 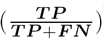 is the fraction of items in *G* that are aligned in *M.* A method with a perfect alignment output would have both precision and recall values of 1.0.

### 5.3 Benchmarking Method

We benchmark HDP-Align against two established alignment methods: SIMA [33] and MZmine2’s Join Aligner [22]. The performance of both methods have been evaluated in past studies [1,14,22,32,33]. SIMA is a stand-alone program while Join aligner is an integral part of the MZmine2 suite widely used for the pre-processing of LC-MS data. The selection of SIMA and Join as the benchmark methods is motivated by the fact that both methods are direct matching methods (thus easily comparable to HDP-Align) but still differ sufficiently in how they establish the final alignment results. This is primarily due to the differences between both methods in the form of the distance/similarity function between peak features, the actual matching algorithm itself and the merging order of pairwise results to construct the full alignment results.

The two most important parameters to configure in both methods are the mass and RT tolerance parameters, used for thresholding and computing feature similarities during matching. We label these common parameters as the *T*_(*m/z*)_ and *T*_*rt*_ parameters. Note that despite the common label, each method may use the parameter values differently during the alignment process. In our experiments, we let *T*_(*m/z*)_ and *T*_*rt*_ vary within reasonable ranges (details in Section 5.4) and report all performance values generated by each combination of the two parameters.

### 5.4 Parameter Optimisations

Tables 3 and 4 describe the parameter ranges of each method during performance evaluation. For HDP-Align (Table 3), we perform the experiments based on our initial choices on the appropriate parameter values. These are almost certainly less than optimal and can be optimised further. The mass cluster standard deviation 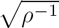 for HDP-ALign is set to the equivalent value in parts-per-million (ppm). These are 500 ppm for the Proteomic dataset and 3 ppm for the Glycomic and Metabolomic datasets. The local (within-run) cluster RT standard deviation 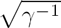 is assumed to be fairly constant and set to 2 seconds for all datasets, while the global cluster standard deviation 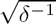 is set in the following dataset-specific manner: 50 seconds for the Proteomic dataset and 20 seconds for the remaining datasets. The larger standard deviation value is required for the Proteomic dataset to accomodate for greater RT drifts across runs. Other hyperparameters in HDP-Align are fixed to the following values: *α'* = 10, *α*_*t*_ = 10, *α*_*m*_ = 100. The values of the precision hyperparameters for global cluster RT (*σ*_0_) and mass cluster (*ρ*_0_) are set to a broad value of 1/5E6. No significant changes were found to the results when these hyperparameters for the DP concentrations and cluster precisions were varied. The mean hyperparameters *μ*_0_ and *ψ*_0_ are set to the means of the RT and m/z values of the input data respectively. During inference, 10000 posterior samples were obtained with the first 5000 used as burn-in, and taking every 10-th sample after burn-in for the posterior probabilities of peaks to be matched.

**Table 3.** Parameters used for HDP-Align

| Dataset | HDP |
| --- | --- |
| P1 Frac 000 | $\sqrt{\rho^{-1}} = 500 \text{ ppm}, \sqrt{\gamma^{-1}} = 2 \text{ s}, \sqrt{\delta^{-1}} = 50 \text{ s}$ |
| P1 Frac 020 |  |
| P1 Frac 040 |  |
| P1 Frac 060 |  |
| P1 Frac 080 |  |
| P1 Frac 100 |  |
| Glycomic | $\sqrt{\rho^{-1}} = 3 \text{ ppm}, \sqrt{\gamma^{-1}} = 2 \text{ s}, \sqrt{\delta^{-1}} = 20 \text{ s}$ |
| Metabolomic | $\sqrt{\rho^{-1}} = 3 \text{ ppm}, \sqrt{\gamma^{-1}} = 2 \text{ s}, \sqrt{\delta^{-1}} = 20 \text{ s}$ |

For SIMA and Join, we report the results from all combinations of the mass and RT tolerance parameters within reasonable ranges listed in Table 4. The ranges of *T*_(*m/z*)_ and *T*_*rt*_ parameters used are based values reported on [14] for the Proteomic dataset and [32] for the Glycomic dataset. For the Metabolomic dataset, they were chosen in light of the mass accuracy and RT deviations of the data.

**Table 4.** Parameters used for the benchmark methods (SIMA, Join).

| Dataset | Benchmark (SIMA, Join) |
| --- | --- |
| P1 Frac 000 | $T_{(m/z)} = \{1.0, 1.1, \dots, 2.0\}, T_{rt} = \{10, 20, \dots, 180\} \text{ s}$ |
| P1 Frac 020 |  |
| P1 Frac 040 |  |
| P1 Frac 060 |  |
| P1 Frac 080 |  |
| P1 Frac 100 |  |
| Glycomic | $T_{(m/z)} = \{0.05, 0.1, 0.25\}, T_{rt} = \{5, 10, \dots, 120\} \text{ s}$ |
| Metabolomic | $T_{(m/z)} = \{0.001, 0.01, 0.1\}, T_{rt} = \{5, 10, \dots, 120\} \text{ s}$ |

In HDP-Align, the mass cluster standard deviation 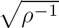 is set to the equivalent value in parts-per-million (ppm). These are 500 ppm for the Proteomic dataset and 3 ppm for the Glycomic and Metabolomic datasets. The local (within-run) cluster RT standard deviation 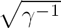 is assumed to be fairly constant and set to 2 seconds for all datasets, while the global cluster standard deviation 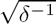 is set in the following dataset-specific manner: 50 seconds for the Proteomic dataset and 20 seconds for the remaining datasets. The larger standard deviation value is required for the Proteomic dataset to accomodate for greater RT drifts across runs.

**Table 5.** Dataset sizes and parameter optimisations.

| Dataset | # runs<br>(total<br>fea-<br>tures) | HDP-Align | SIMA, Join |
| --- | --- | --- | --- |
| **Proteomic** |  |  |  |
| Frac 000 | 2 (10606) | $\sqrt{\rho^{-1}} = 500$<br>ppm,<br>$\sqrt{\gamma^{-1}} = 2$ s,<br>$\sqrt{\delta^{-1}} = 50$ s | $T_{(m/z)} =$<br>$\{1.0, 1.1, \dots, 2.0\},$<br>$T_{rt} =$<br>$\{10, 20, \dots, 180\}$<br>s |
| Frac 020 | 2 (2135) |  |  |
| Frac 040 | 2 (2188) |  |  |
| Frac 060 | 2 (3342) |  |  |
| Frac 080 | 2 (2086) |  |  |
| Frac 100 | 2 (1326) |  |  |
| **Glycomic** | 10 (9344) | $\sqrt{\rho^{-1}} = 3$<br>ppm,<br>$\sqrt{\gamma^{-1}} = 2$ s,<br>$\sqrt{\delta^{-1}} = 20$ s | $T_{(m/z)} =$<br>$\{0.05, 0.1, 0.25\},$<br>$T_{rt} =$<br>$\{5, 10, \dots, 120\}$ s |
| **Metabolomic** | 6 (7477) | $\sqrt{\rho^{-1}} = 3$<br>ppm,<br>$\sqrt{\gamma^{-1}} = 2$ s,<br>$\sqrt{\delta^{-1}} = 20$ s | $T_{(m/z)} =$<br>$\{0.001, 0.01, 0.1\},$<br>$T_{rt} =$<br>$\{5, 10, \dots, 120\}$ s |

Other hyperparameters in HDP-Align are fixed to the following values: *α*′ = 10, *α*_*t*_ = 10, *α*_*m*_ = 100. The values of the precision hyperparameters for global cluster RT (*σ*_0_) and mass cluster (*ρ*_0_) are set to a broad value of 1/5E6. No significant changes were found to the results when these hyperparameters for the DP concentrations and cluster precisions were varied. The mean hyperparameters *μ*_0_ and *ψ*_0_ are set to the means of the RT and m/z values of the input data respectively. During inference for the Glycomic and Metabolomic datasets, 500 posterior samples were collected for the Gibbs sampling procedure, discarding the first 500 during the burn-in period. For the Proteomic dataset with larger RT deviations, 5000 posterior samples were obtained after discarding the first 5000 samples during burn-in. The number of samples is selected to ensure convergence during inference.

## 6 Results

Precision and recall values for the evaluated methods methods on the different datasets are shown in Sections 6.1 and 6.2. Additionally, an example of the further annotations for the putative adduct type and metabolite identity that can be produced by HDP-Align is also shown in Section 6.2. Running time of the evaluated methods are reported in Section 6.3.

### 6.1 Proteomic Results

Figure 4 shows the results from performance evaluation on the Proteomic dataset. We see that both benchmark methods (SIMA and Join) produce a wide range of performance depending on the parameter values for (*T*_(*m/z*)_,*T*_*rt*_) chosen. Sensitivity to parameter values is expected on this dataset due to the low mass accuracy in the MS instrument that produces the data and the high RT drifts present across runs (further details in [14]). HDP-Align performs well on several fractions (particularly fractions 040, 060, 080, 100) with precision-recall performance close to the optimal performance attainable by the benchmark methods. On all fractions, HDP-Align is also able to produce higher-precision results compared to the benchmark methods by reducing recall through setting the appropriate values for the threshold *t*. The primary benefits of quantifying alignment uncertainties is realised here as the well-calibrated probability scores on the matching confidence of aligned peak features produced HDP-Align allows the user to choose which point along the PR curve to operate on. It is less obvious how this can be accomplished in the benchmark methods by varying the RT (*T*_*rt*_) and m/z (*T*_*m/z*_) thresholding parameters, if at all possible.

**Figure 4.**
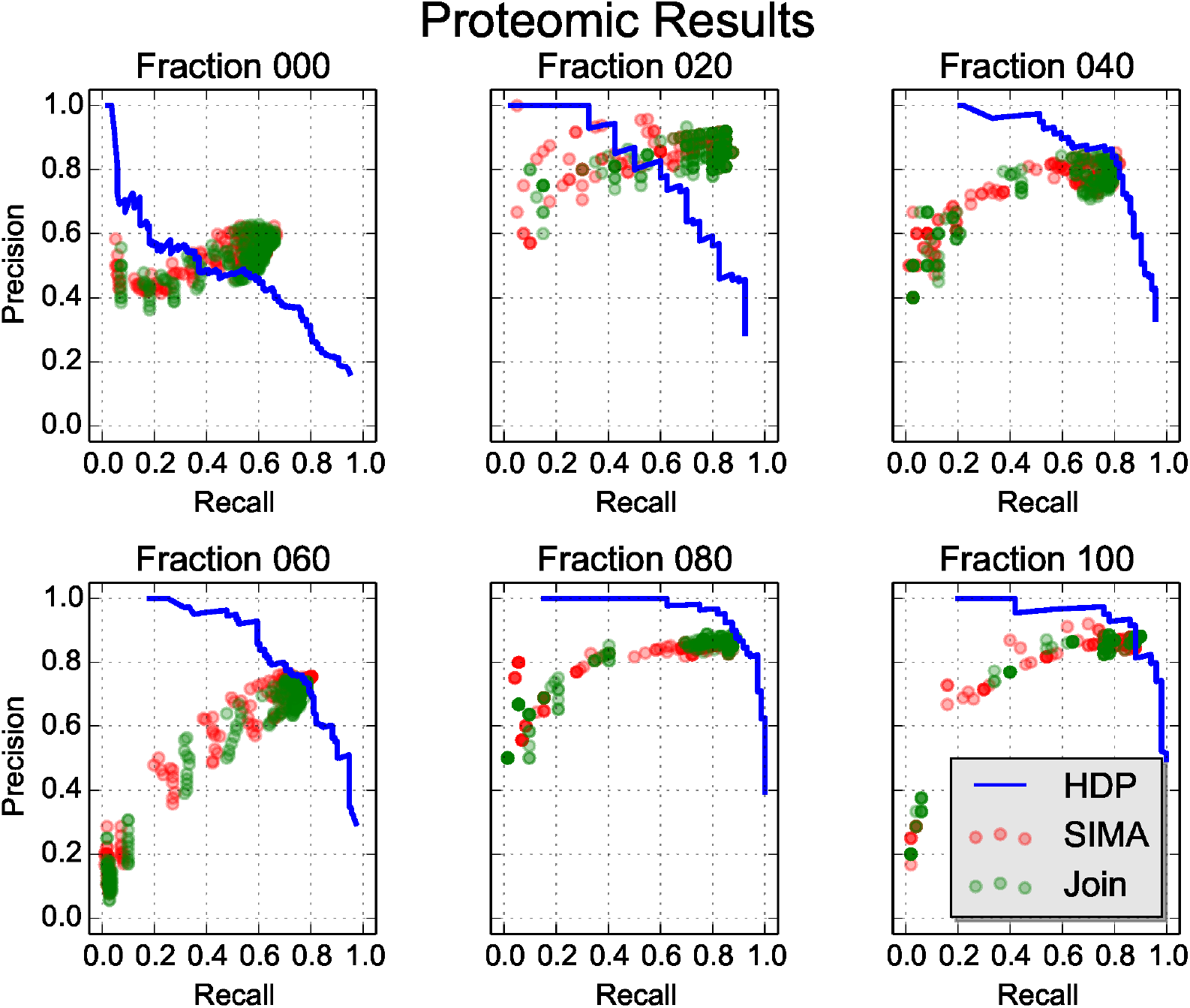
Precision-recall values on the different fractions of the Proteomic dataset.

### 6.2 Glycomic and Metabolomic Results

Figures 5 and 6 show the results from experiments on the Glycomic and Metabolomic datasets. Similar to the Proteomic dataset, a wide range of precision-recall values can be observed in the results for the benchmark methods on the two datasets. The performance of HDP-Align, using the same set of parameters on both datasets, come close to the optimal results from the benchmark methods, while still allowing the user to control the desired point along the precision-recall curve to operate on.

**Figure 5.**
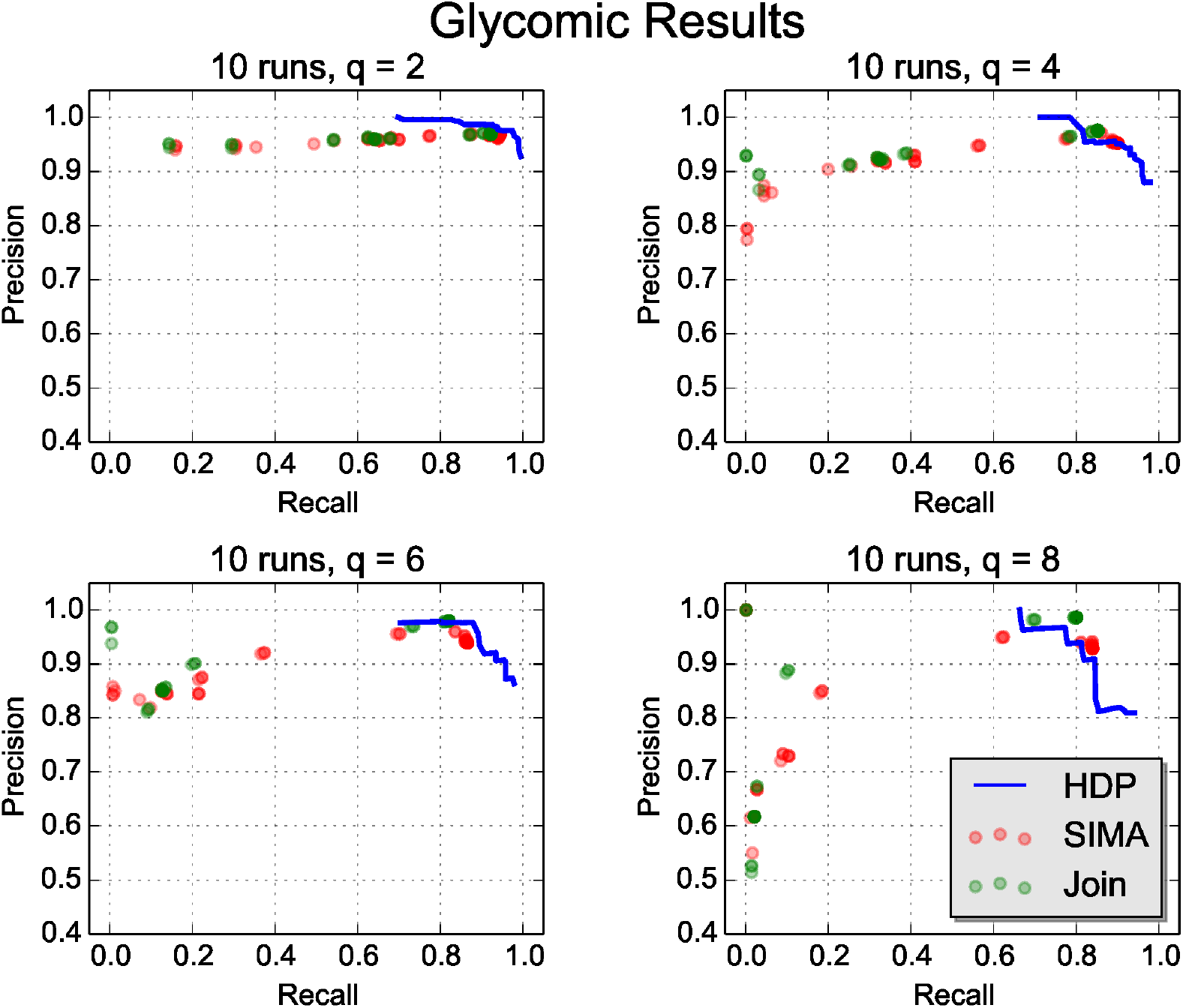
Precision-recall values on the alignment of 10 runs from the Glycomic dataset when *q* (the strictness of performance evaluation as described in Section 5.2) is gradually increased.

**Figure 6.**
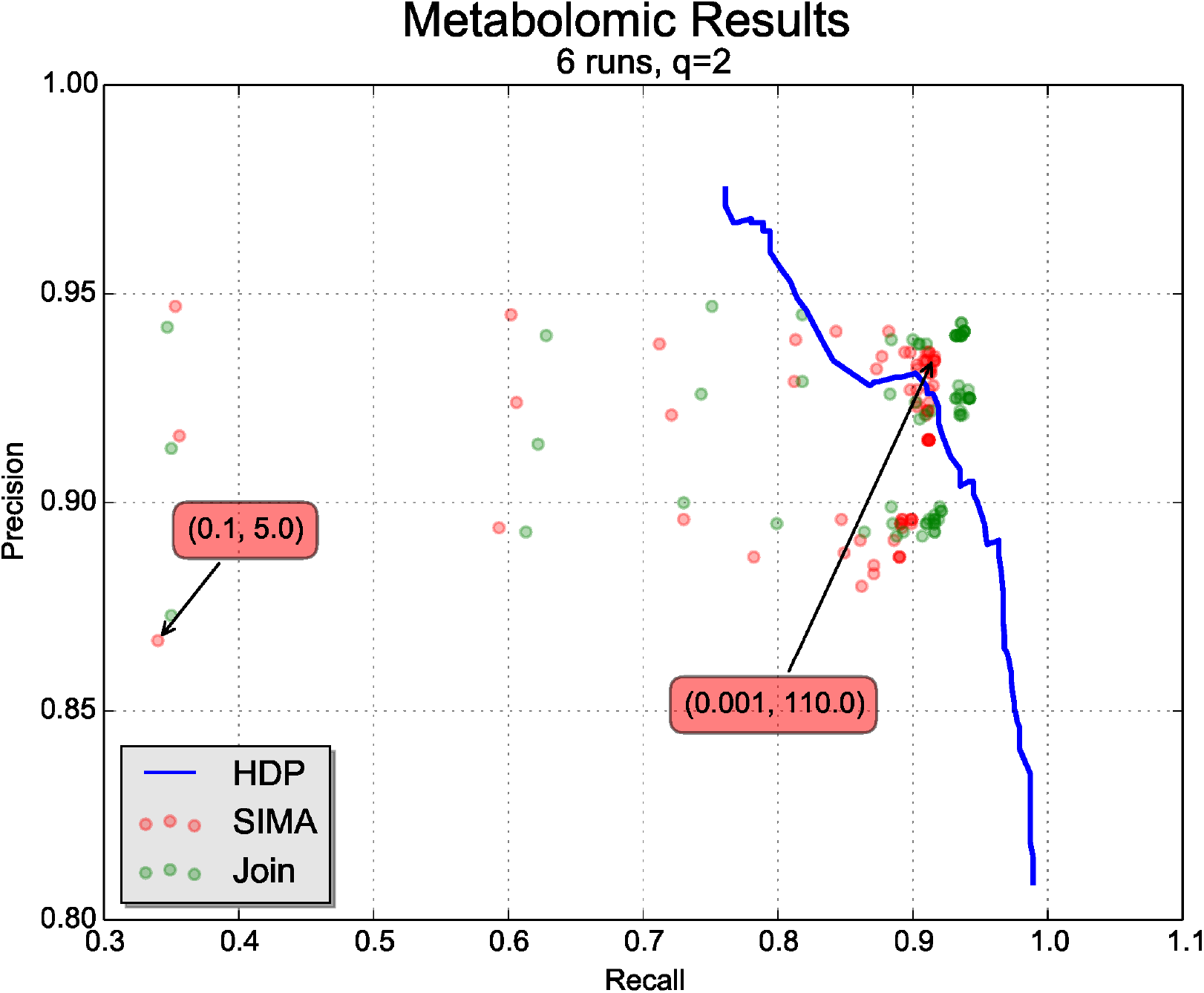
Precision-recall values on the alignment of 6 runs from the Metabolomic dataset. The parameter values (*T*_*m/z*_,*T*_*rt*_) that produce the best and worst performance in SIMA are also annotated in the Figure (red boxes).

The results for the Glycomic dataset (Figure 5) also show some additional results on how the measured precision-recall values might change depending on the strictness of what constitutes an ‘item’ during performance evaluation. This is accomplished by gradually increasing the value for *q* (described in detail in Section 5.2) that determines the size of the feature combinations enumerated from a method’s output. For example, *q*=2 considers all pairwise combinations of features from the method’s output during performance evaluation, while *q* = 4 considers all combinations of size 4, and so on. Figure 5 shows that as *q* is increased, parameter sensitivity seems to become more of an issue for the benchmark methods, with more parameter sets having lower precisions in the results. Across all *q*s evaluated, parameter pairs that produce the best alignment performance (points with high precision and recall values) are generally small *T*_(*m/z*)_ and large *T*_*rt*_ values. Examples of parameter pairs that produce the best and worse performance for SIMA are shown in Figure 6. The results here appear to suggest the importance of having high mass precision during matching. Importantly, we see from Figure 5 that the performance of HDP-Align remains fairly consistent as *q* is increased.

The Metabolomic dataset also provides us with additional results in form of annotations of putative adduct type and metabolite identities. A thorough evaluation on the quality of such annotations, in comparison to e.g. the workflow proposed in [15], is beyond the scope of this paper and would likely necessitate using a different and more apppopriate evaluation dataset. Instead, we present an example of the further analysis performed by HDP-Align (as proposed in Section 4.4) on the resulting clustering objects after inference. Figure 7 shows a global RT cluster where peak features across runs have been grouped by their RT and m/z values. Within this global cluster, peak features are further separated into 6 mass clusters – corresponding to ionisation products produced by the global cluster during mass spectometry. In Figure 7, mass cluster *A* and *B* contain features aligned from several runs but they do not have any other mass cluster sharing a possible precursor mass. Mass cluster *C* and *D* share a common precursor mass (292.12696) and can thus be annotated by the adduct type that produce the transformation. Similarly, mass cluster *E* and *F* share a common precursor mass at 383.14278. Queries to a local KEGG database are issued based on the precursor mass values, producing several compound identities that can be putatively assigned to the global RT cluster. It is a great strength of our approach that this putative identification step appears very naturally from the alignment results.

**Figure 7.**
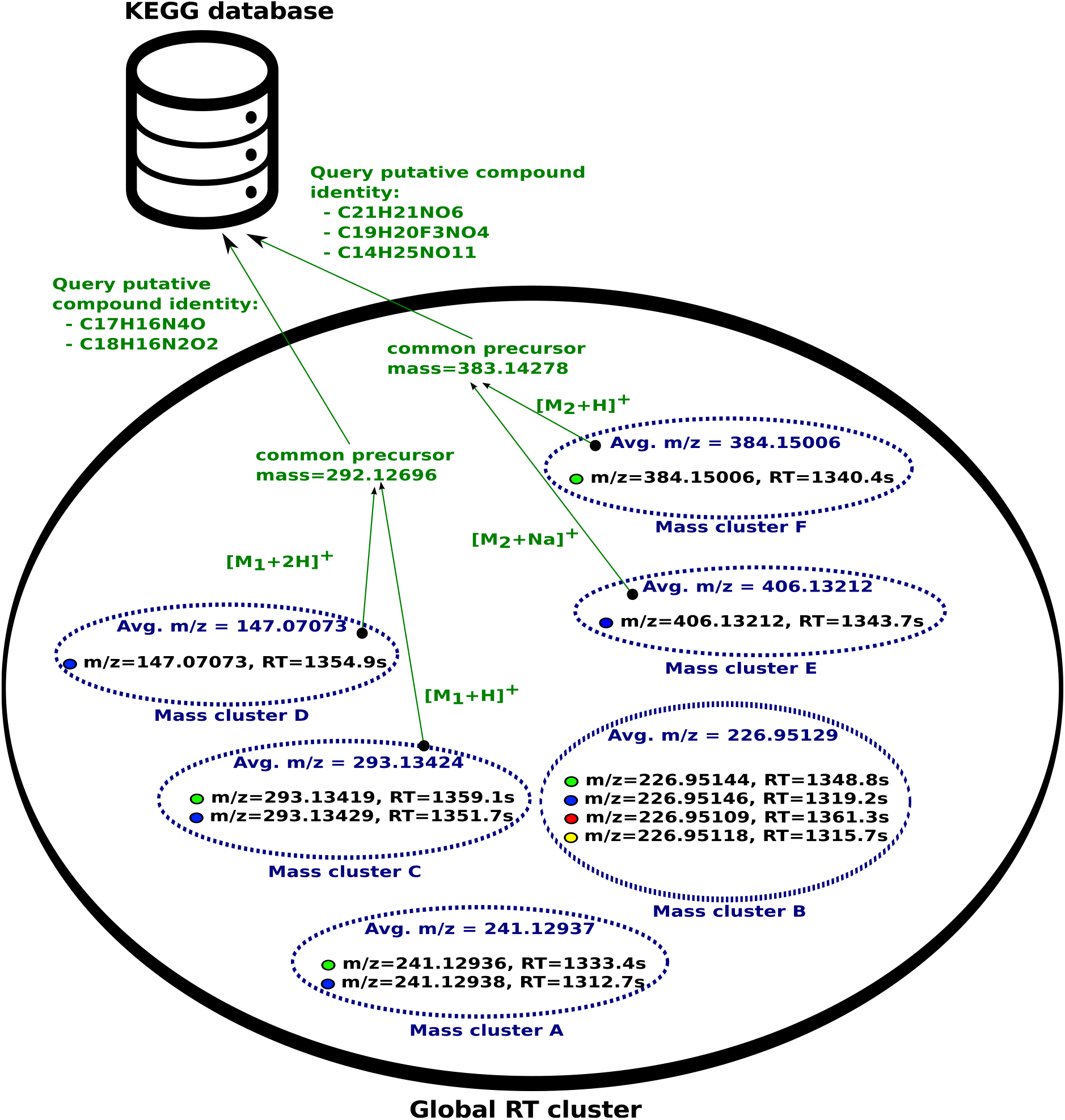
Example analysis that can be performed on the clustering objects inferred in a Gibbs sample from HDP-Align. The outer black oval denotes a global RT cluster (generally corresponding to a metabolite compound), while the smaller dotted ovals within denote mass clusters (labelled as mass cluster *A*, *B*, *C*, *D*, *E*, *F*). Peak features are denoted by the filled circle, with the fill colour indicating the originating run of a peak. Green colours denote additional analysis steps that can be performed on the mass cluster objects.

### 6.3 Running Time

The main factor affecting the running time of HDP-Align is the total number of peaks across all runs to be processed and the number of samples produced during Gibbs sampling. In each iteration of Gibbs sampling, HDP-Align removes a peak from the model, updates parameters of the model conditioned on every other parameters, and reassigns a peak into RT and mass clusters. The time complexity of this operation is *O*(*N*), where *N* is the total number of peaks across all runs. In practice, additional time will also be spent on various necessary book-keeping operations, such as deleting empty clusters that are no longer required, updating internal data structures, etc.

A representative running time is given as *N* = 9344 for the Glycomic dataset. HDP-Align requires approximately 5 hours to collect 1000 samples. In comparison, both SIMA and Join perform alignment within 5 to 10 seconds. Similarly, for *N* = 7477 for the Metabolomic dataset, HDP-Align produces the results in approximately 4 hours after collecting 1000 samples, while SIMA and Join complete within seconds. The running time of HDP-Align, while being significantly longer than these two benchmark methods, is comparable to other computationally-intensive steps (e.g. peak detection) in a typical LC-MS pipeline.

## 7 DISCUSSION AND CONCLUSION

We have presented a hierarchical non-parametric Bayesian model that performs direct matching of peak features, a problem of significant importance in the data pre-processing pipeline of large untargeted LC-MS datasets. Unlike other direct matching methods, the novelty of our proposed approach lies in its ability of to produce well-calibrated probability scores on the matching confidence of aligned peak features (evidenced by the increasing precision and decreasing recall as the threshold *t* is increased). This is accomplished by casting the multiple alignment problem of LC-MS peak features as a hierarchical clustering problem. Matching confidence can then be obtained based on the probabilities of co-eluting peak features to be assigned under the same mass component of the same global cluster. Experiments based on datasets from real proteomic, glycomic and metabolomic experiments show that HDP-Align is able to produce alignment results competitive to the benchmark alignment methods, with the added benefit of being able to provide a measure of confidence in the alignment quality. This can be useful in real analytical situations, where neither the optimal parameters nor the alignment ground truth is known to the user.

Through comparisons against benchmark methods, our studies have also investigated the effect of sub-optimal parameter choices on alignment performance. While beyond the scope of our paper, we agree with [27, 28] that thorough investigations into the influence of numerous configurable parameters (prevalent in nearly all LC-MS data processing pipeline) on the resulting biological conclusions are of utmost importance. This should be followed by the development of methods to minimise or automatically-tune such configurable parameters. Despite the abundance of new methods proposed for LC-MS data pre-processing, relatively few studies have been done on the subject of quantifying uncertainties and alleviating the burden of parameter optimisations during actual data analysis. One way to minimise the number of parameters is through the integration of multiple steps in the typical LC-MS pipeline into fewer steps. Our proposed model in HDP-Align can potentially be extended in this manner, as evidenced by the metabolomic dataset results where we directly use the clustering objects inferred from the model to perform further analysis on putative adduct and metabolite type annotations. While the proposed annotation approach in Section 4.4 is fairly simple, it can be easily extended to more sophisticated annotation strategies, such as in CAMERA [12].

A primary weakness of HDP-Align lies in the long computational time required to produce results. Additional work will be required to reduce the computational burden of the model through various optimisation tricks and potentially by parallelising the Gibbs inference step using e.g. the method described in [17]. Another possibility is to adopt a different non-sampling-based inferential approach while still retaining the essence and benefits of the HDP model in this paper. The results presented in the current paper suggest the method shows enough promise to warrant the effort to speed it up.

Another aspect worthy of investigation is determining the most effective way to present and visualise the alignment probabilities produced by HDP-Align. Additional sources of information present in the LC-MS data, such as chromatographic peak shapes, can also be used to improve alignment performance and subsequent analyses that follow.

Finally, replacing or enhancing the mixture of mass components used in HDP-Align with a more appropriate mass model, such as that in MetAssign [4] that specifically takes into account the inter-dependency structure of peaks, is an avenue for future work. This will be particularly useful when extending the proposed model in HDP-Align into a single inferential model that encompasses many intermediate steps in a typical LC-MS data processing pipeline.

## Acknowledgments

We would like to thank Glasgow Polyomics for providing us with the metabolomics data used for performance evaluation. JW was funded by a PhD studentship from SICSA. SR was supported by the BBSRC (BB/L018616/1).

## References

1. R. Ballardini, M. Benevento, G. Arrigoni, L. Pattini, and A. Roda. MassUntangler: A novel alignment tool for label-free liquid chromatography–mass spectrometry proteomic data. Journal of Chromatography A, 1218(49):8859–8868, 2011.

2. R. K. Bradley, A. Roberts, M. Smoot, S. Juvekar, J. Do, C. Dewey, I. Holmes, and L. Pachter. Fast statistical alignment. PLoS Comput. Biol., 5(5):e1000392, May 2009.

3. D. J. Creek, A. Jankevics, R. Breitling, D. G. Watson, M. P. Barrett, and K. E. V. Burgess. Toward global metabolomics analysis with hydrophilic interaction liquid chromatography–mass spectrometry: improved metabolite identification by retention time prediction. Analytical Chemistry, 83(22):8703–8710, 2011.

4. R. Daly, S. Rogers, J. Wandy, A. Jankevics, K. E. Burgess, and R. Breitling. Metassign: probabilistic annotation of metabolites from LC-MS data using a Bayesian clustering approach. Bioinformatics, 30(19):2764–2771, 2014.

5. D. P. De Souza, E. C. Saunders, M. J. McConville, and V. a. Likić. Progressive peak clustering in GC-MS Metabolomic experiments applied to Leishmania parasites. Bioinformatics, 22(11):1391–6, June 2006.

6. R. C. H. De Vos, S. Moco, A. Lommen, J. J. B. Keurentjes, R. J. Bino, and R. D. Hall. Untargeted large-scale plant metabolomics using liquid chromatography coupled to mass spectrometry. Nat. Protoc., 2(4):778–91, Jan. 2007.

7. B. Fischer, J. Grossmann, V. Roth, W. Gruissem, S. Baginsky, and J. M. Buhmann. Semi-supervised LC/MS alignment for differential proteomics. Bioinformatics, 22(14):e132–40, July 2006.

8. M. Ghanat Bari, X. Ma, and J. Zhang. PeakLink: a new peptide peak linking method in LC-MS/MS using wavelet and SVM. Bioinformatics, 30(17):2464–70, Sept. 2014.

9. A. Jankevics, M. E. Merlo, M. de Vries, R. J. Vonk, E. Takano, and R. Breitling. Separating the wheat from the chaff: a prioritisation pipeline for the analysis of metabolomics datasets. Metabolomics, 8(1):29–36, 2012.

10. J. Jeong, X. Shi, X. Zhang, S. Kim, and C. Shen. Model-based peak alignment of metabolomic profiling from comprehensive two-dimensional gas chromatography mass spectrometry. BMC Bioinformatics, 13(1):27, Jan. 2012.

11. X. Kong and C. Reilly. A Bayesian approach to the alignment of mass spectra. Bioinformatics, 25(24):3213–20, Dec. 2009.

12. C. Kuhl, R. Tautenhahn, C. Böttcher, T. R. Larson, and S. Neumann. CAMERA: an integrated strategy for compound spectra extraction and annotation of liquid chromatography/mass spectrometry data sets. Anal. Chem., 84(1):283–9, Jan. 2012.

13. G. Landan and D. Graur. Characterization of pairwise and multiple sequence alignment errors. Gene, 441(1-2):141–7, July 2009.

14. E. Lange, R. Tautenhahn, S. Neumann, and C. Gröpl. Critical assessment of alignment procedures for LC-MS proteomics and metabolomics measurements. BMC Bioinformatics, 9:375, 2008.

15. T. S. Lee, Y. S. Ho, H. C. Yeo, J. P. Y. Lin, and D.-Y. Lee. Precursor mass prediction by clustering ionization products in LC-MS-based metabolomics. Metabolomics, 9(6):1301–1310, Apr. 2013.

16. J. Listgarten, R. M. Neal, S. T. Roweis, and A. Emili. Multiple alignment of continuous time series. In Advances in neural information processing systems, pages 817–824, 2004.

17. D. Lovell, R. P. Adams, and V. Mansingka. Parallel markov chain monte carlo for dirichlet process mixtures. In Workshop on Big Learning, NIPS, 2012.

18. C. D. Manning, P. Raghavan, and H. Schütze. Introduction to information retrieval, volume 1. Cambridge University Press, Cambridge, 2008.

19. C. Notredame, D. G. Higgins, and J. Heringa. T-Coffee: A novel method for fast and accurate multiple sequence alignment. J. Mol. Biol., 302(1):205–17, Sept. 2000.

20. O. Penn, E. Privman, G. Landan, D. Graur, and T. Pupko. An alignment confidence score capturing robustness to guide tree uncertainty. Mol. Biol. Evol., 27(8):1759–67, Aug. 2010.

21. V. Perera, M. D. T. Zabala, H. Florance, N. Smirnoff, M. Grant, and Z. R. Yang. Aligning extracted lc-ms peak lists via density maximization. Metabolomics, 8(1):175–185, 2012.

22. T. Pluskal, S. Castillo, A. Villar-Briones, and M. Orešič. MZmine 2: Modular framework for processing, visualizing, and analyzing mass spectrometry-based molecular profile data. BMC Bioinformatics, 11(1):395, 2010.

23. K. Podwojski, A. Fritsch, D. C. Chamrad, W. Paul, B. Sitek, K. Stühler, P. Mutzel, C. Stephan, H. E. Meyer, et al. Retention time alignment algorithms for LC/MS data must consider non-linear shifts. Bioinformatics, 25(6):758–764, 2009.

24. B. D. Redelings and M. a. Suchard. Joint Bayesian estimation of alignment and phylogeny. Syst. Biol., 54(3):401–18, June 2005.

25. R. a. Scheltema, A. Jankevics, R. C. Jansen, M. a. Swertz, and R. Breitling. PeakML/mzMatch: a file format, Java library, R library, and tool-chain for mass spectrometry data analysis. Anal. Chem., 83(7):2786–93, 2011.

26. C. a. Smith, E. J. Want, G. O’Maille, R. Abagyan, and G. Siuzdak. XCMS: processing mass spectrometry data for metabolite profiling using nonlinear peak alignment, matching, and identification. Analytical chemistry, 78(3):779–87, Feb. 2006.

27. R. Smith, D. Ventura, and J. T. Prince. LC-MS alignment in theory and practice: a comprehensive algorithmic review. Briefings in Bioinformatics, 2013. DOI:10.1093/bib/bbt080.

28. R. Smith, D. Ventura, and J. T. Prince. Novel algorithms and the benefits of comparative validation. Bioinformatics, 29(12):1583–5, June 2013.

29. Y. W. Teh, M. I. Jordan, M. J. Beal, and D. M. Blei. Hierarchical dirichlet processes. Journal of the American Statistical Association, 101(476):1566–1581, 2006.

30. J. D. Thompson, D. G. Higgins, and T. J. Gibson. CLUSTAL W: improving the sensitivity of progressive multiple sequence alignment through sequence weighting, position-specific gap penalties and weight matrix choice. Nucleic Acids Res., 22(22):4673–80, Nov. 1994.

31. R. Tibshirani, T. Hastie, B. Narasimhan, S. Soltys, G. Shi, A. Koong, and Q.-T. Le. Sample classification from protein mass spectrometry, by ‘peak probability contrasts’. Bioinformatics, 20(17):3034–3044, 2004.

32. T.-H. Tsai, M. G. Tadesse, C. Di Poto, L. K. Pannell, Y. Mechref, Y. Wang, and H. W. Ressom. Multi-profile Bayesian alignment model for LC-MS data analysis with integration of internal standards. Bioinformatics, 29(21):2774–2780, 2013.

33. B. Voss, M. Hanselmann, B. Y. Renard, M. S. Lindner, U. Köthe, M. Kirchner, and F. A. Hamprecht. SIMA: Simultaneous multiple alignment of LC/MS peak lists. Bioinformatics, 27(7):987–993, 2011.

